# Spiny mice (*Acomys*) exhibit attenuated hallmarks of aging and rapid cell turnover after UV exposure in the skin epidermis

**DOI:** 10.1101/2020.05.07.083287

**Authors:** Wesley Wong, Austin Kim, Ashley W. Seifert, Malcolm Maden, Justin D. Crane

## Abstract

The study of long-lived and regenerative animal models has revealed diverse protective responses to stressors such as aging and tissue injury. Spiny mice (*Acomys*) are a unique mammalian model of skin regeneration, but their response to other types of physiological skin damage have not been investigated. In this study, we examine how spiny mice skin responds to acute UVB damage or chronological aging compared to non-regenerative C57Bl/6 mice (*M. musculus)*. We find that, compared to *M. musculus*, the skin epidermis in *A. cahirinus* experiences a similar UVB-induced increase in basal cell proliferation but exhibits increased epidermal turnover. Notably, *A. cahirinus* uniquely form a suprabasal layer co-expressing Keratin 14 and Keratin 10 after UVB exposure concomitant with reduced epidermal inflammatory signaling and reduced markers of DNA damage. In the context of aging, old *M. musculus* animals exhibit typical hallmarks including epidermal thinning, increased inflammatory signaling and senescence. However, these age-related changes are absent in old *A. cahirinus* skin. Overall, we find that *A. cahirinus* have evolved novel responses to skin damage that reveals new aspects of its regenerative phenotype.

## Introduction

Maintenance and repair of the skin barrier are essential to prevent infection and protect the body from external hazards. However, normal stress and damage during an organism’s lifespan deteriorates skin structure and repair through undefined mechanisms. The skin epidermis is composed primarily of keratinocytes derived from basal stem cells that continually renew the differentiated spinous, granular, and cornified suprabasal layers (Fuchs, 2008). Together, these cells protect against environmental insults such as radiation, physical injury, dehydration, and pathogens. A major environmental hazard to the skin is ultraviolet (UV) solar radiation, which induces potentially mutagenic DNA damage if left unrepaired. UV-damaged cells undergo either apoptosis or cell cycle arrest coupled with DNA repair before re-entry into the cell cycle which results in thickening of the epidermis (El-Abaseri et al., 2006). Aging of the skin induces cellular senescence, thinning of the epidermis and reduced basal stem cell renewal and proliferation (Liu et al., 2019). While we are beginning to understand some of the mechanisms that underlie the slow repair of aged skin (Keyes et al., 2016; Wong et al., 2019), further research is needed.

Long-lived or stress-resistant model organisms permit a better understanding of adaptive mechanisms of tissue homeostasis and repair. For example, the Snell dwarf mouse attains a long lifespan through impaired pituitary hormone signaling and its study has informed our understanding of how organismal physiology and longevity are balanced (Brown-Borg, 2011; Flurkey et al., 2002). The naked mole rat *Heterocephalus glaber* (*H. glaber)* has evolved distinctive mechanisms of cancer resistance which may be mediated by microenvironment and immune system (Hadi et al., 2018; Seluanov et al., 2009; Tian et al., 2015). In a similar vein, models of vertebrate regeneration such as salamanders and newts have been used extensively to understand the optimal healing of nervous, muscle and connective tissue (Joven et al., 2019), but a critical barrier to deriving clinical therapies from this work has been a lack of mammalian models of tissue repair. The recent discovery that spiny mice (*Acomys spp.,* hereafter referred to as *Acomys*) can fully regenerate skin and hair following wounding or burn injury (Maden, 2018; Seifert et al., 2012), a feature not observed in typical strains of mice (*Mus musculus; Mus*), lends well to uncovering novel regenerative mechanisms in skin tissue. However, it is not known how *Acomys* responds to other types of skin stress, including UV-damage and chronological aging.

Here, we examine epidermal responses to acute UVB exposure and chronological aging in the conventional C57Bl/6 *M. musculus* strain compared to *A. cahirinus*. We find that *A. cahirinus* has attenuated pathological hallmarks of UVB exposure and aging in conjunction with distinctive differences in epidermal turnover, senescence and inflammation. These processes may underlie the unique skin regeneration phenotype of *A. cahirinus*, and such mechanisms can inform therapeutic strategies to manage skin diseases such as skin cancer and aging.

## Materials and Methods

### Mice

For all studies, C57Bl/6 (*M. musculus)* mice were acquired from Jackson Labs (#000664). For the UVB experiments, 5-6 month old male *A. cahirinus* animals were shipped from the University of Florida and compared to 3-4 month old male C57Bl/6 mice. *M. musculus* were housed 4-5 to a cage in mouse cages while *A. cahirinus* were housed 2-3 to a cage in rat cages with a 12h/12h light cycle. Animals were housed at Northeastern University for at least 2 weeks prior to UVB or sham treatments to acclimate. For the aging studies, young, female animals of each species at 3-4 months of age were used and old, female animals (*M. musculus*: 2 years of age*; A. cahirinus*: 4 years of age) were analyzed. The young and old *M. musculus* mice were housed at Northeastern University while the young and old *A. cahirinus* animals were housed at the University of Florida and University of Kentucky. For the UVB studies, all animals were sacrificed by cervical dislocation. For the aging studies, animals were euthanized either by cervical dislocation (*M. musculus*) or isoflurane overdose (*A. cahirinus*). All procedures were performed according to IACUC approved procedures from each institution.

### Ultraviolet Irradiation

Animals were sedated with intraperitoneal ketamine/xylazine anesthesia (50 mg/kg ketamine; 5 mg/kg xylazine) and shaved with clippers prior to UVB exposure. Animals were exposed to a UVB dose of 200 mJ/cm^2^ using a dosimeter calibrated UV instrument (310 nm peak output) (Tyler Research, Alberta, Canada). Full-thickness dorsal skin samples in telogen phase were collected and fixed at 24- and 48-hours after irradiation in addition to sham controls and paraffin embedded cross-sections were prepared for histologic analysis. BrdU (5-Bromo-2’-doxyuridine, Sigma Aldrich, St. Louis, Missouri, USA) was injected intraperitoneally at 100 mg/kg in sterile PBS 24 hours prior to sacrifice. Animals were sacrificed by cervical dislocation.

### Histology

Tissues were fixed overnight in 10% neutral buffered formalin and thereafter transferred to 70% ethanol at 4°C until processing. Tissues were processed using an automated tissue processor (Thermo HistoStar, Kalamazoo, Michigan, USA) and embedded in paraffin wax. A microtome (Leica Biosystems, Buffalo Grove, Illinois, USA) was used to cut 4 μm skin cross-sections, which were allowed to dry overnight before de-waxing and further processing. Heat induced epitope retrieval was performed on de-waxed slides with either citrate or Tris-EDTA buffer prior to immunofluorescence staining. Skin cross-sections were blocked for 30 minutes at room temperature in 5% normal goat serum PBS, which was also the primary antibody diluent, unless the target was a phospho-protein in which case 5% normal goat serum TBS was used. Primary antibodies and their usage were as follows: Keratin 14 (chicken, BioLegend #906004, 1:500, San Diego, California, USA), HMGB1 (rabbit, Abcam #AB79823, 1:250, Cambridge, Massachusetts, USA), Lamin B1 (rabbit, Abcam #AB16048, 1:1000), Keratin 10 (rabbit, BioLegend #905404, 1:500), Loricrin (rabbit, BioLegend #905104, 1:500), BrdU (rat, Abcam #AB6326, 1:200), thymine dimer mouse, Novus #NB600-1141, 1:400, Centennial, Colorado, USA), cleaved caspase-3 (rabbit, Cell Signaling #9661, 1:100, Danvers, Massachusetts, USA), γH2AX (rabbit, Cell Signaling #9718, 1:40). Primary antibody incubations were carried out overnight at 4°C except Keratin 14, Keratin 10, and Loricrin, which were 2 hours at room temperature. Following either PBS or Tris-buffered saline (TBS) wash, secondary detection antibodies conjugated with either AlexaFluor-647 or AlexaFluor-488 (Invitrogen, Carlsbad, California, USA) were diluted in either PBS or TBS at 1:200 were applied for 30 minutes at room temperature. Mounting media with DAPI (4’,6-diamidino-2-phenylindole) used was either ProLong Gold (Invitrogen) or ProLong Diamond (Invitrogen), which were used interchangeably and allowed to cure prior to imaging. For thymine dimer staining, a mouse-on-mouse blocking kit (Vector Labs #BMK-2202, Burlingame, California, USA) was used per the manufacturer’s instructions.

Hematoxylin and Eosin staining was performed on 4 um de-waxed sections using standard histology protocols using Gill’s Hematoxylin No.1 and Eosin Y (Sigma Aldrich). Colorimetric stained slides were mounted with Permount (Fisher Scientific, Pittsburgh, Philadelphia, USA). All brightfield imaging was done using an inverted EVOS XL (Life Technologies, Carlsbad, California) microscope. All fluorescence imaging was performed on either an Revolve R4 (Echo Labs, San Diego, California, USA) microscope equipped with Olympus UPlanFL 10x/0.30, and UPlanFL 20x/0.50 air objectives using DAPI, FITC, and Cy5 fluorescence channels or an Axio Observer Z1 (Zeiss, White Plains, New York, USA), equipped with Plan Apochromat 5x/0.16, 10x/0.45, and 20x/0.8 air objectives and 40x/1.3 and 63x/1.4 oil objectives using DAPI, GFP, DsRed, and Cy5 fluorescence channels.

### Image analysis

All image processing and analysis was conducted using ImageJ FIJI (version 2.0.0-rc-69/1.52i, NIH), Photoshop CC 2019 (version 20.0.4, Adobe), and Illustrator CC 2019 (version 23.0.3, Adobe, San Jose, California, USA). Immunofluorescence images first underwent thresholding over entire image set of all samples from the experiment followed by manual counting of positive cells for at least 3 unique fields from each sample. All epidermal analyses only counted interfollicular epidermal cells and excluded hair follicles. Nuclear HMGB1 positive cells were those that had overlapping signal with their nuclear DAPI signal. Lamin B1 positive cells were those that exhibited perinuclear signal relative to their DAPI signal. Keratin 14, Keratin 10, and Loricrin positive cells were those that had DAPI-associated cytoplasmic signal.

Epidermal thickness measurements were performed on H&E stained intact skin cross-sections as previously described (Crane et al., 2015) by measuring the orthogonal distance from the basement membrane to the outer edge of the cellular epidermis.

### Gene Expression

RNA was extracted by homogenizing whole skin in TRIzol (Invitrogen) reagent using a bead mill apparatus (MPBio 5G, Irvine, California, USA) followed by column purification (Zymo Direct-zol RNA mini prep, Irvine, California, USA) with on column DNase treatment and subsequent elution. 600 ng of total RNA was then reversed transcribed to cDNA (ABI HC cDNA synthesis kit, Waltham, Massachusetts, USA), diluted 1:30 in ultrapure water and mRNA expression was assessed using qPCR with SYBR chemistry. CDK-interacting protein 1 (*Cdkn1a*) was used as a housekeeping gene and its expression did not significantly differ between species or with UVB treatment. Data was analyzed by the delta Ct (ΔCt) method.

Primers are as follows:

*M. musculus Cxcl1*-Forward: ACTCAAGAATGGTCGCGAGG; *M. musculus Cxcl1*-Reverse: GTGCCATCAGAGCAGTCTGT. *A. cahirinus Cxcl1*-Forward: CCCATGGTTCGGAAGGTTGT; *A. cahirinus Cxcl1*-Reverse: GTTGTCAGACGCCAGCATTC. *M. musculus Il1b*-Forward: TGCCACCTTTTGACAGTGATG; *M. musculus Il1b*-Reverse: TGATGTGCTGCTGCGAGATT. *A. cahirinus Il1b*-Forward: CTGGGCTCCAGAGACACAAG; *A. cahirinus Il1b*-Reverse: GAACCCCTGCATCAACTCCA. *M. musculus Cdkn1a*-Forward: CGGTGTCAGAGTCTAGGGGA; *M. musculus Cdkn1a*-Reverse: AGGATTGGACATGGTGCCTG. *A. cahirinus Cdkn1a*-Forward: TGCACTCTGGTATCTCACGC; *A. cahirinus Cdkn1a*-Reverse: CAGTCGGCGCTTAGAGTGAT.

### Statistical Analysis

All data were analyzed in Prism 8.0 (GraphPad, San Diego, California, USA) using a two-way analysis of variance (ANOVA) on either log or square root transformed data with Tukey’s HSD post-hoc testing for differences between treatments and species. Statistical significance was set as *p<0.05*.

## Results

### UV-irradiation induced changes to skin epidermal morphology and epidermal stratification of *M. musculus* and *A. cahirinus*

While *A. cahirinus* can fully regenerate skin wounds and burns without scarring (Maden, 2018; Seifert et al., 2012), it has not been established how this regenerative ability translates to other types of skin damage. Since UVB radiation is a common form of epidermal damage (Brash et al., 1991), we sought to compare the morphological changes between *A. cahirinus* and *M. musculus* epidermis following an acute dose of UVB known to induce epidermal proliferation and DNA damage in *M. musculus* (Trevithick et al., 1992). We first measured the total thickness of the cellular epidermis using hematoxylin and eosin staining. As expected, *M. musculus* had a significantly thicker epidermis at 48 hours post-UVB compared to sham or 24 hours (Fig. 1A). In contrast, there was no appreciable difference in epidermal thickness in response to UVB in *A. cahirinus* at 24 hours (*p=0.21*) or 48 hours (*p=0.80*) compared to sham (Fig. 1B). To understand the processes underlying this differential response, we examined rates of cell division by analyzing post-UVB epidermal incorporation of the synthetic nucleoside 5-bromo-2’-deoxyuridine (BrdU). During imaging, we noted species specific patterns of BrdU labeling in basal and suprabasal epidermal compartments (Fig. 1C). To distinguish these changes, we analyzed patterns of label retention in the two compartments separately. In the basal epidermis, BrdU incorporation was significantly lower in *M. musculus* basal epidermis 24 hours after UVB-exposure, and significantly higher at 48 hours following UVB compared to both sham and 24 hours (Fig 1D). In contrast, *A. cahirinus* epidermis did not exhibit changes in BrdU incorporation at 24 hours post-exposure but was significantly elevated at 48 hours compared to both the sham and 24 hour groups (Fig. 1D). In the suprabasal epidermis, both *M. musculus* and *A. cahirinus* exhibited greater BrdU incorporation at 48 hours compared to sham, however this UV-induced proliferation was greater in *M. musculus* (Fig. 1E).

**Figure 1.**
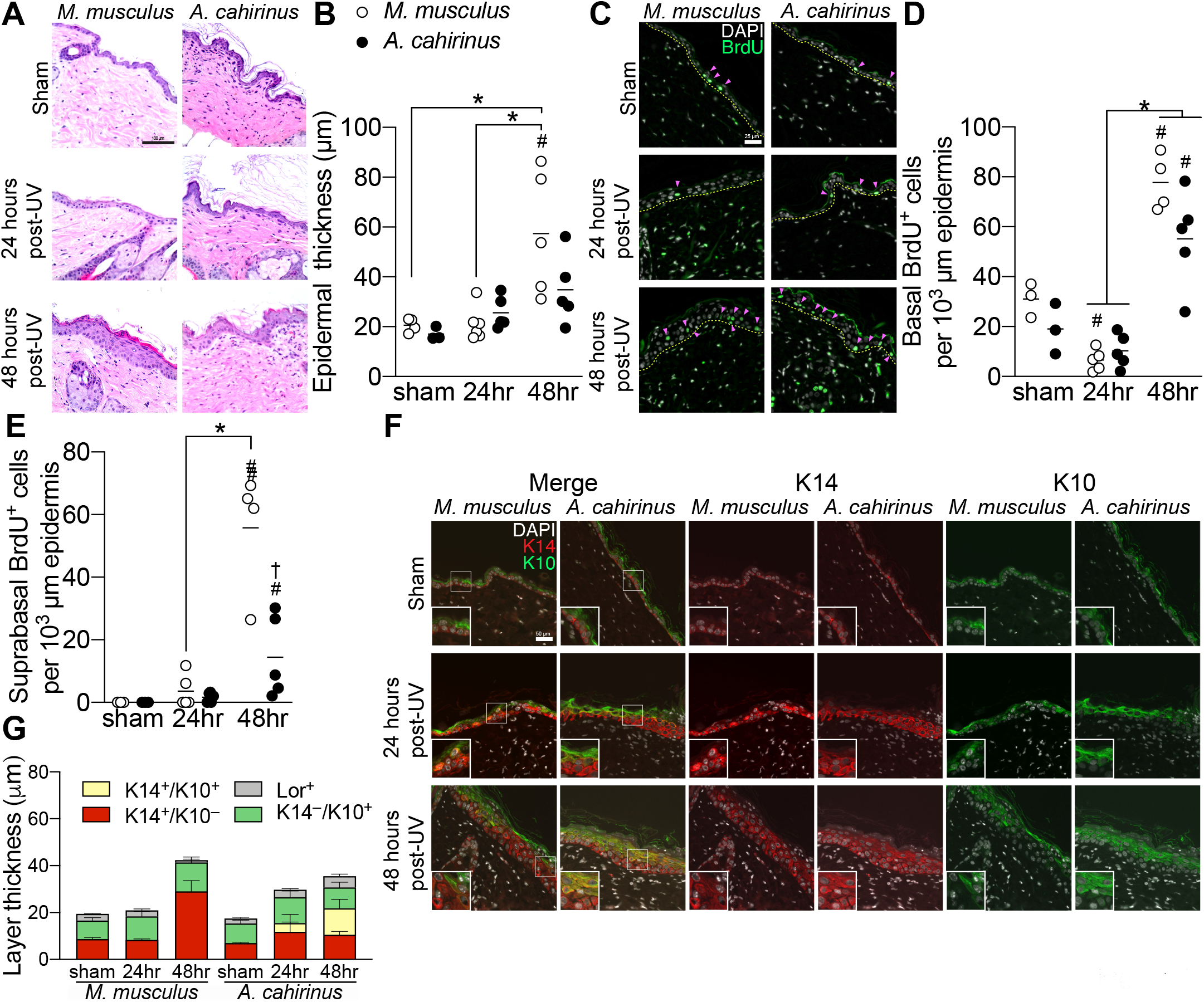
Acute UVB-exposure induces distinctive pattern of skin epidermal differentiation in *A. cahirinus*. *A*, representative brightfield microscopy images of hematoxylin and eosin-stained skin from control (*sham*) and UV-irradiated *M. musculus* and *A. cahirinus*, collected 24 and 48 hours after exposure. Scale bar = 100 μm. *B*, quantification of epidermal thickness. *n = sham both speces, 3; 24hr M. musculus, 6; 24hr A. cahirinus, 5; 48hr both species, 5 mice per group*. *C*, representative immunofluorescence images of epidermal BrdU labeling. The epidermal basement membrane is indicated by the *yellow dashed line*. Positive cells are indicated by the pink arrows. *Scale bar = 25 μm. D and E*, quantification BrdU labeling in (*D*) basal epidermis and (*E*) suprabasal epidermis. *n = sham both species, 3; 24hr both species, 5; 48hr M. musculus, 4; 48hr A. cahirinus, 5 mice per group*. *F*, representative immunofluorescence images of epidermal differentiation markers keratin 14 (*K14*) and keratin 10 (*K10*) labeling with magnified inset of the basal-suprabasal junction. *Scale bar = 50 μm*. *G*, quantification of individual differentiation marker layer thickness. *n = 3 mice per group*. *Loricrin images and individual layer comparisons included in Supplemental Figure 1A-E.* Data points are biological replicates. Lines indicate group means. *, significantly different from the indicated group (*p<0.05*). ^#^, significantly different relative to sham control (*p<0.05*). ^†^, significantly different (*p<0.05)* to *M. musculus* at the same timepoint.

Since stem cell fate decisions control the balance of cell division and upward transport through the suprabasal layers (Williams et al., 2011), we next stained for keratin 14 (K14) to mark the basal stem/progenitor layer, keratin 10 (K10) to mark the spinous layer, and for the Loricrin (Lor) expressing cornified envelope after UVB in each species. During imaging, we noticed species specific patterns of UVB-induced epidermal differentiation, most notably the unique double-positive K14^+^/K10^+^ suprabasal layer in UVB-exposed *A. cahirinus* (Fig. 1F). We then measured the thicknesses of each of the labeled basal (K14^+^/K10^−^), double-positive (K14^+^/K10^+^), spinous (K14^−^/K10^+^), and cornified (Lor^+^) epidermal layers. Exposure to acute UVB resulted in a significantly thicker K14^+^ layer in *M. musculus* by 48 hours but not in *A. cahirinus*. However, a double-positive K14^+^/K10^+^ suprabasal epidermal layer was uniquely evident only in *A. cahirinus* at 24 and 48 hours post-UVB (Fig. 1G). The thickness of the Lor^+^ layer between species was similar at all time points and was not statistically different (*p=0.0986)* in *A. cahirinus* at 48 hours (Fig. 1G and Fig. S1A). Overall, *A. cahirinus* skin exhibits a unique program of epidermal stratification without hyperproliferation. This distinctive pattern of differentiation consists of a transient intermediate layer with both basal and spinous characteristics in response to UVB exposure.

### Differences in UV-induced damage and epidermal cell death between *M. musculus* and *A. cahirinus*

In *M. musculus* and humans, differentiation and eventual shedding of skin cells via upward transport through the epithelium serves as a method of eliminating damaged and compromised cells (Freije et al., 2014). Since we observed an altered differentiation pattern in *A. cahirinus* in response to UVB, we reasoned that this could impact the removal of damaged cells. We first assessed this by measuring the UVB-induced DNA photoproduct thymine dimer (T-T dimer) in the epidermis (Fig. 2A). As expected from this dose of UVB, we found significantly higher thymine dimer positive cells in *M. musculus* basal epidermis by 24 hours with a near return to sham levels by 48 hours (Fig. 2B), yet in the suprabasal layer levels rose significantly by 24 hours and remained elevated at 48 hours (Fig. 2C). In *A. cahirinus*, while we found an induction of thymine dimer positive cells 24 hours following UVB, this UV-response was reduced compared to *M. musculus* animals in both the basal and suprabasal epidermis (Fig. 2B-C). To extend our thymine dimer results, we next assessed the DNA repair marker γH2AX (Fig. 2D). *M. musculus had greater* basal and suprabasal epidermal γH2AX labeling at 24 hours and 48 hours after UVB (Fig. 2E and 2F). *A. cahirinus* also had more γH2AX-positive cells versus sham at 24 hours in the basal layer, with a similar pattern in the suprabasal layers at 24 and 48 hours (Fig. 2E and 2F). However, similar to thymine dimers, there were more γH2AX-positive cells in *M. musculus* compared to *A. cahirinus* at 24 and 48 hours post UVB exposure (Fig. 2E and 2F).

**Figure 2.**
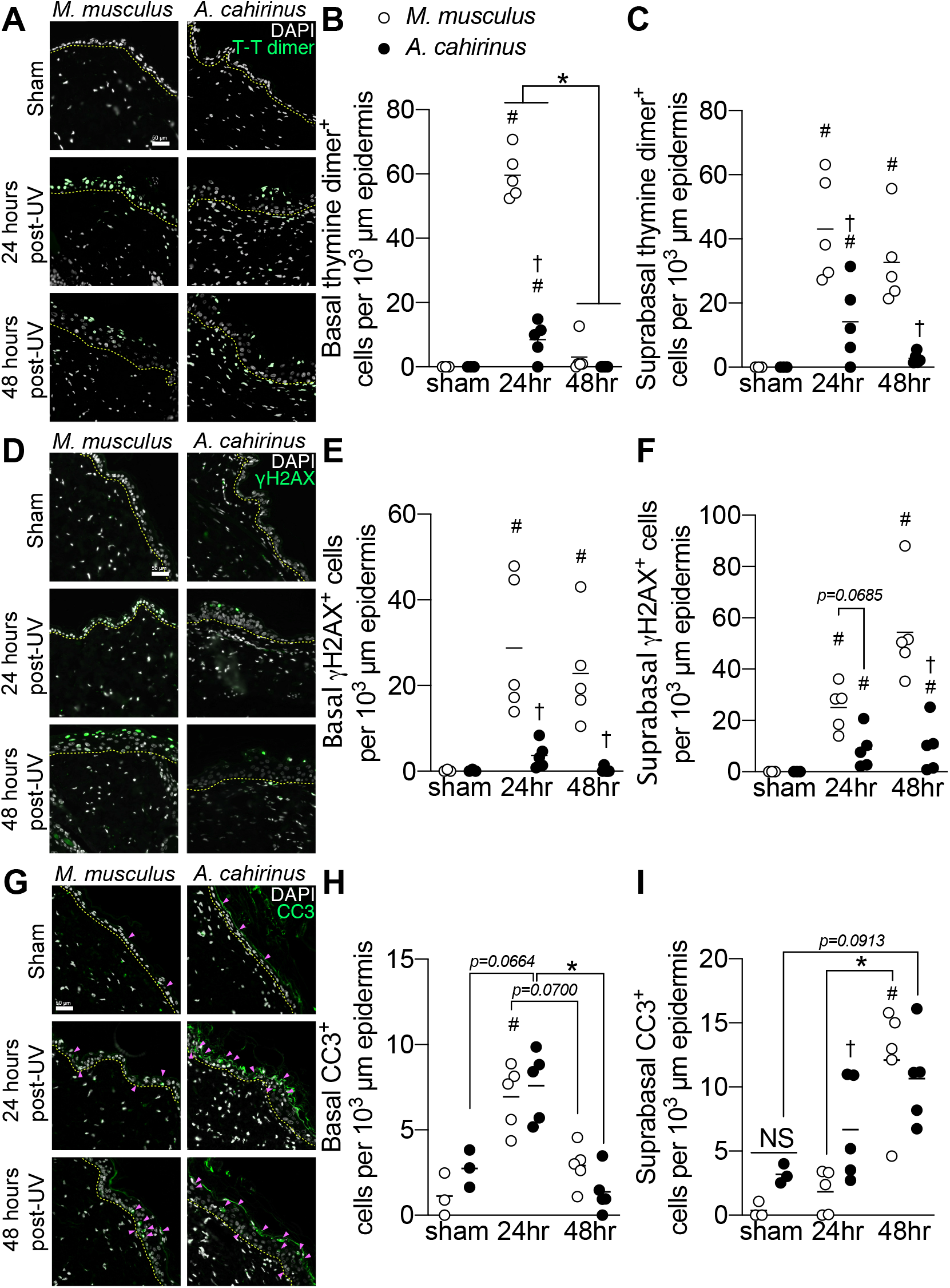
Efficient removal of damaged and dying skin epidermal cells through rapid turnover and apoptosis in acute UVB-exposed *A. cahirinus*. *A*, representative immunofluorescence images of epidermal thymine dimer (*T-T dimer*) labeling of skin from control (*sham*) and UV-irradiated *M. musculus* and *A. cahirinus*, collected 24 and 48 hours after exposure. *B and C*, quantification of thymine dimer labeling in (*B*) basal epidermis and (*C*) suprabasal epidermis. *D*, representative immunofluorescence images of epidermal γH2AX labeling*. E and F*, quantification γH2AX labeling in (*D*) basal epidermis and (*E*) suprabasal epidermis. *G*, representative immunofluorescence images of epidermal cleaved caspase-3 (*CC3)* labeling. Positive cells are indicated by the pink arrows. *H and I*, quantification of cleaved caspase-3 labeling in (*H*) basal and (*I)* suprabasal epidermis. Data points are biological replicates. Lines indicate group means. *, significantly different from the indicated group (*p<0.05*). ^#^, significantly different relative to sham control (*p<0.05*). ^†^, significantly different (*p<0.05)* to *M. musculus* at the same timepoint. The epidermal basement membrane is indicated by the *yellow dashed line. All scale bars = 50 μm. All measurements n = sham both speces, 3; 24hr both species, 5; 48hr both species, 5 mice per group*.

To understand the contribution of apoptosis to the post-UVB epidermal response, we measured the abundance of cleaved caspase-3 (CC3) positive cells (Fig. 2G). In the basal layer, there was a similar UVB-induced response between species, with an increase in cleaved caspase-3 expressing cells at 24 hours followed by a return to sham levels 48 hours later (Fig. 2H). This was in contrast to the suprabasal epidermis, where at 24 hours *A. cahirinus* exhibited a significantly greater number of cleaved caspase-3 expressing cells compared to *M. musculus. However, there* was a similar abundance of cleaved caspase-3 expressing cells between species at 48 hours (*M. musculus* significantly elevated *vs* baseline; *p=0.0913* for *A. cahrinus vs* baseline) (Fig. 2I). Thus, *A. cahirinus* epidermis have modest changes in cell death responses following UVB exposure compared to *M. musculus*.

### Attenuated skin epidermal inflammatory response following UV-irradiation in *A. cahirinus*

Inflammation is a major component of epidermal remodeling following UVB exposure (Barker et al., 1991). To gain further insight into the differential UVB responses between *M. musculus* and *A. cahirinus*, we analyzed the epidermal abundance of the damage associated molecular pattern, HMGB1 (Fig. 3A). Loss of nuclear HMGB1 is a biomarker of cell stress, senescence, inflammation, and autophagic responses particularly in UVB-exposed basal epidermis (Bald et al., 2014; Davalos et al., 2013). Quantification of HMGB1 labeling revealed that *M. musculus* experiences a significant loss of basal nuclear HMGB1 by 48 hours after UVB (Fig. 3B). However, we found that post-UVB *A. cahirinus* basal epidermis retained nuclear HMGB1 labeling similar to sham levels (Fig. 3B). Suprabasal levels of HMGB1 were similar between species and treatments (Fig. 3C), suggesting that this effect is restricted to basal progenitors. A milieu of pro-inflammatory signaling within the skin also occurs concomitantly to the epidermal hyperplastic response (Ouhtit et al., 2000). To better understand the UV-induced inflammatory response in each species, we measured mRNA levels of the inflammation associated genes *Il1b*, *Cxcl1, Tgfb1, and Mmp9* in whole skin using species specific primers and qPCR. Increased epidermal expression of *Il1b* and *Cxcl1* mediate inflammatory responses after UV-exposure (Qiang et al., 2017), and knockout mouse studies have shown loss of *Tgfb1* and *Mmp9* are associated with increased and prolonged inflammation in the skin epidermis (Wang et al., 1999; Yoshinaga et al., 2008). Acute UVB-irradiation resulted in significant increase in *Cxcl1* in *M. musculus* skin at 48 hours, but levels in *A. cahirinus* were not significantly altered by UVB exposure (Fig. 3D). In contrast, *Il1b* expression was not significantly changed by acute UVB-irradiation in either species but was consistently lower in *A. cahirinus* vs. *M. musculus* (Fig. 3E). Transcript levels of *Tgf1b* in *M. musculus* skin significantly decreased at 24 hours but *A. cahirinus* levels were unchanged (Fig. 3F). Similarly, levels of *Mmp9* in *M. musculus* skin significantly decreased at 24 and 48 hours but *A. cahirinus* levels were not significantly altered by UVB (Fig. 3G). Thus, compared to *M. musculus, A. cahirinus* epidermis resists UVB-driven inflammatory gene expression changes, consistent with their anti-inflammatory tissue damage response seen after wounding and burn injury (Brant et al., 2015; Gawriluk et al., 2019; Maden, 2018).

**Figure 3.**
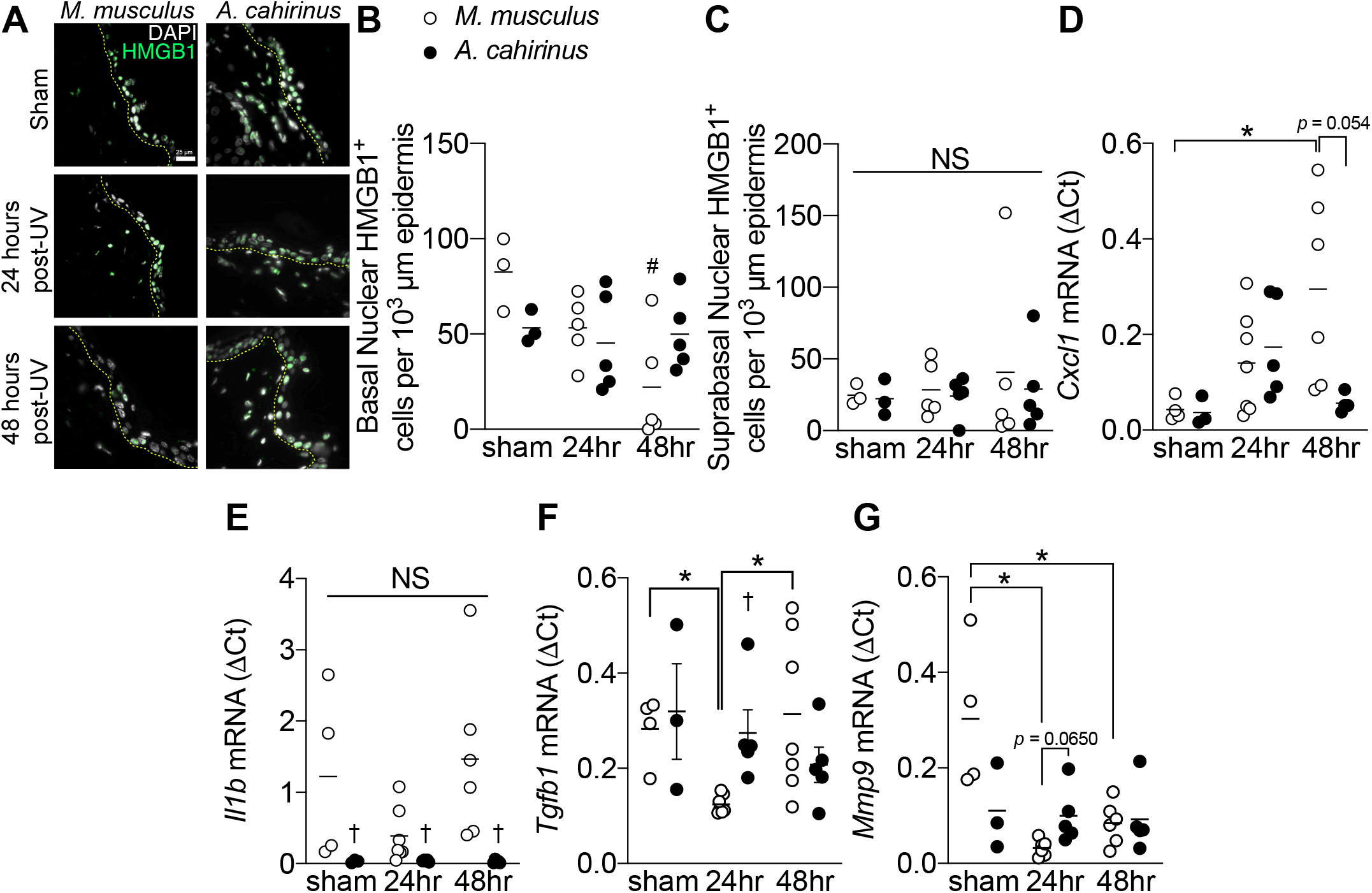
Attenuated skin epidermal inflammatory response following UV-irradiation in *A. cahirinus*. *A*, representative immunofluorescence images of epidermal HMGB1 labeling of skin from control (*sham*) and UV-irradiated *M. musculus* and *A. cahirinus*, collected 24 and 48 hours after exposure. The epidermal basement membrane is indicated by the *yellow dashed line. Scale bar = 50 μm. B and C*, quantification of nuclear HMGB1 labeling in (*B*) basal epidermis and (*C*) suprabasal epidermis. *n = sham both speces, 3; 24hr both species, 5; 48hr both species, 5 mice per group. D*, *E, F, and G*, mRNA expression (ΔCt) of (*D) Cxcl1*, (*E*) *Il1b*, (*F*) *Tgfb1*, and (*G*) *Mmp9* in each treatment group as measured by qPCR*. n = sham M. musculus, 3; sham A. cahirinus, 3; 24hr M. musculus, 6, 24hr A. cahirinus, 5; 48hr M. musculus, 6; 48hr A. cahirinus, 3 mice per group for Cxcl1*. *n = sham M. musculus, 4; sham A. cahirinus, 3; 24hr M. musculus, 6; 24hr A. cahirinus, 3; 48hr M. musculus, 5; 48hr A. cahirinus, 3 mice per group for Il1b*. *n = sham M. musculus, 4; sham A. cahirinus, 3; 24hr M. musculus, 5; 24hr A. cahirinus, 5; 48hr M. musculus, 6; 48hr A. cahirinus, 5 mice per group for Tgfb1*. *n = sham M. musculus, 4; sham A. cahirinus, 3; 24hr M. musculus, 5; 24hr A. cahirinus, 5; 48hr M. musculus, 6; 48hr A. cahirinus, 5 mice per group for Mmp9.* Data points are biological replicates. Lines indicate group means. *, significantly different from the indicated group (*p<0.05*). ^#^, significantly different relative to sham control (*p<0.05*). ^†^, significantly different (*p<0.05)* to *M. musculus* at the same timepoint.

### Aging associated epidermal thinning, inflammatory signaling, and senescence are absent in *A. cahirinus*

Total organismal aging is the sum of chronological aging and environmental exposure. We sought to further explore the differences in cellular stress response in these two species by examining their epidermal responses to chronological aging. *A. cahirinus* have a maximal lifespan of 5.9 years, versus 3.5 years in *M. musculus* (C57Bl/6) (Edrey et al., 2012). Thus, to study aging in these species, we used young animals from each species at 3-4 months old as well as aged 2-year old *M. musculus* and 4-year old *A. cahirinus* animals. At these ages the species are roughly matched as a fraction of their respective maximum lifespans: approximately 9.4% for young *M. musculus*, 8.4% for young *A. cahirinus*, 56% for old *M. musculus*, and 68% for old *A. cahirinus.* Using formalin fixed skin cross-sections for both species, we used hematoxylin and eosin staining of the skin to reveal morphological changes due to age (Fig. 4A). This revealed an expected decrease in the thickness of the cellular epidermis of old versus young *M. musculus* (Fig. 4B), however, the cellular epidermal thickness of *A. cahirinus* remained unchanged with age (Fig. 4B). To better understand the cellular epidermal thickness differences, we measured the nuclear envelope protein and senescence associated biomarker Lamin B1 (Fig. 4C). Similar to previous research by ourselves (Wong et al., 2019) and others (Freund et al., 2012; Wang et al., 2017), we found a decrease in basal epidermal Lamin B1 with age in *M. musculus* (Fig. 4D). However, *A. cahirinus* basal epidermal Lamin B1 labeling was similar to young *M. musculus* and was not different in old *A. cahirinus* versus young (Fig. 4D). To corroborate these findings regarding senescence, we also evaluated the proportion of epidermal cells with nuclear HMGB1 staining (Fig. 4E). As previously reported by ourselves (Wong et al., 2019) and others (Davalos et al., 2013), we observed a decrease in epidermal nuclear HMGB1^+^ cells with age in *M. musculus* (Fig. 4F), yet HMGB1 in old *A. cahirinus* remained unchanged from young animals (Fig. 4F). Collectively, these data show that *A. cahirinus* have reduced hallmarks of epidermal aging.

**Figure 4.**
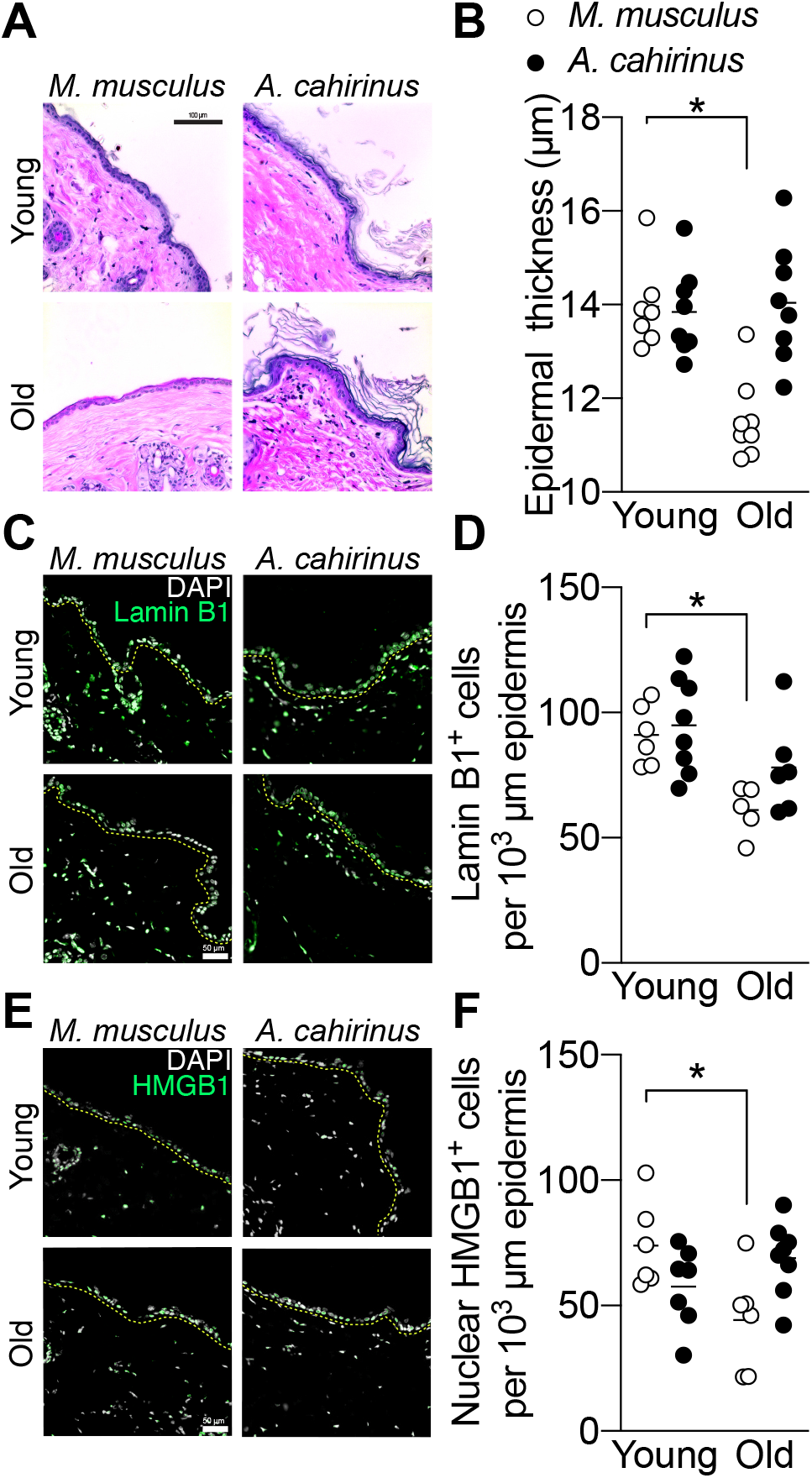
Aging associated epidermal thinning, inflammation, and senescence are absent in *A. cahirinus*. *A*, representative brightfield microscopy images of hematoxylin and eosin-stained intact skin from control (*Young)* and aged (*Old) M. musculus* and *A. cahirinus. Scale bar = 100 μm. B*, quantification of epidermal thickness. *C,* representative immunofluorescence images of epidermal Lamin B1 labeling of skin from each treatment group. The epidermal basement membrane is indicated by the *yellow dashed line. Scale bar = 50 μm. n = young M. musculus, 7; young A. cahirinus, 8; old both species, 8 mice per group. D*, quantification of epidermal Lamin B1 labeling*. n = young M. musculus, 6; young A. cahirinus, 8; old M. musculus, 5, old A. cahirinus, 6 mice per group. E*, representative immunofluorescence images of epidermal HMGB1 labeling of skin from each treatment group. The epidermal basement membrane is indicated by the *yellow dashed line. Scale bar = 50 μm. F*, quantification of epidermal nuclear HMGB1 labeling. *n = young M. musculus, 6; young A. cahirinus, 7; old M. musculus, 6, old A. cahirinus, 7 mice per group.* Data points are biological replicates. Lines indicate group means. *, significantly different from the indicated group (*p<0.05*).

## Discussion

The comparative study of exceptionally long-lived, stress-resistant, or regenerative organisms can be used to provide insight into evolved mechanisms of cellular repair. In this study, we found substantial differences in how the regenerative rodent *A. cahirinus* responds to UVB skin damage compared to *M. musculus*. This included altered patterns of keratinocyte proliferation and differentiation, an earlier induction of UVB-induced cell death, and an attenuated inflammatory response. The acanthosis commonly observed in *M. musculus* after UVB was not evident in *A. cahirinus* despite similar rates of basal cell proliferation, suggesting an alternative fate of newly formed keratinocytes. Notably, the *A. cahirinus* epidermis forms a unique middle suprabasal layer of keratinocytes with both basal and spinous characteristics (K14^+^/K10^+^) which may facilitate increased keratinocyte upward transport and removal. This was supported by the observations of fewer retained damaged cells (thymine dimers, γH2AX) as well as more apoptotic cells transiting the suprabasal epidermis at intermediate stages of repair (24 hours) in *A. cahirinus* compared to *M. musculus.* Taken together, these data suggest that *A. cahirinus skin* epidermis is capable of more rapid removal of damaged and dying cells after UVB-exposure through an enhanced rate of differentiation.

The presence of a co-expressed K14^+^/K10^+^ suprabasal layer in the epidermis has been reported by others, but usually this is f in the context of skin disorders. For example, in psoriatic human skin biopsies, there is aberrant Notch expression and dual K14/K10 expression (Ota et al., 2014). Dysregulated epidermal calcium gradients can also produce dual expression of K14 and K10, such as in the skin blistering disorders Hailey-Hailey disease and Darier disease (Mikkelsen et al., 2018; Robia and Young, 2018), and in TRPV4 KO mice (Moore et al., 2013) which suffer defective tight junction formation (Sokabe et al., 2010). Interestingly, TRPV4 is also a nociceptor and those mice had altered pain perception and inflammatory responses after sunburn (Moore et al., 2013). Aberrant cell cycle regulation can also cause dual K14/K10 expression as mice lacking epidermal C/EBPβ and C/EBPα exhibit defective cornified envelope formation (Lopez et al., 2009). In contrast to these pathological examples of K14/K10 co-expression, this suprabasal layer formed after UVB in *A. cahirinus* appears to be a physiological skin damage response. Notably, *A. cahirinus* executes this distinctive program of epidermal differentiation without marked changes in inflammatory signaling factors, which is similar to the blunted inflammatory response reported during wound healing (Brant et al., 2015).

The study of organisms with long lifespans such as the naked mole rat (*H. glaber*) has revealed several unique mechanisms of stress resistance (Pérez et al., 2009; Seluanov et al., 2009). Long-lived model organisms often also exhibit altered inflammatory and cellular damage responses distinct from humans or laboratory mice which provide cancer resistance or augmented tissue repair (Eming et al., 2009; Kowalczyk et al., 2020). In *H. glaber*, cellular senescence is rare, yet cancer rates are very low due to sensitive cellular growth inhibition, modified tumor suppressor pathways, altered extracellular matrix composition, and a stable epigenome (Seluanov et al., 2018). The evolutionary path to these adaptive differences was likely guided by evolutionary adaptation to the hypoxic subterranean environment of *H. glaber* and the fructose biased metabolism resulting from hypoxia (Seluanov et al., 2018). We reasoned that *A. cahirinus* might have also evolved distinct cellular stress mechanisms to enable extensive tissue regeneration, which parallels the unique survival mechanisms recently found in *A. cahirinus* fibroblasts *in vitro* (Saxena et al., 2019). We found that *A. cahirinus* epidermis was remarkably resilient to chronological aging stress with minimal changes in epidermal thickness or biomarkers of aging and senescence. It is interesting to note that the MRL/MpJ healer strains of mice also exhibit reduced inflammatory responses to injury and improved recovery outcomes in the corneal epithelium (Ueno et al., 2005) and ear punch wounds (Kench et al., 1999). Thus, an attenuated inflammatory response may enable the unique growth and repair patterns of *A. cahirinus* epidermis compared to *M. musculus* and may also explain the differences in biological aging between the species.

Our study of the regenerative African spiny mouse, *A. cahirinus*, has revealed a unique skin response to acute UVB-irradiation which consists of rapid differentiation and apoptosis without prototypical epidermal hyperplasia or inflammation. However, we do not yet understand the precise cellular mechanisms underlying this response. While proliferation and differentiation may be intrinsically regulated, inflammation and clearance of cells is likely related to the altered immunity and inflammation previously reported during skin repair in *A. cahirinus* (Maden, 2018; Simkin et al., 2017). Moreover, our findings of improved epidermal tissue repair in *A. cahirinus* extend recent work using models of spinal cord injury (Streeter et al., 2019), musculoskeletal regeneration (Gawriluk et al., 2016), and muscle damage (Maden et al., 2018). Future work should clarify the molecular underpinnings of improved tissue repair in *A. cahirinus* in order to advance regenerative medicine.

## Acknowledgements

The authors wish to thank J. Monaghan for helpful discussions related to the project and E. Crane for assistance with some of the IHC analysis.

## Author contributions

Conceptualization: W.W., J.D.C.; Investigation: W.W., A.K.; Formal Analysis: W.W., A.K.; Resources: J.D.C., A.W.S., M.M.; Writing – original draft: W.W., J.D.C.; Writing – review & editing: W.W., A.K., J.D.C, A.W.S., M.M.; Visualization: W.W.; Supervision: W.W., J.D.C.; Project administration: J.D.C.; Funding acquisition: J.D.C.

## List of abbreviations

K10: Keratin 10
K14: Keratin 14
Lor: Loricrin
PBS: Phosphate buffered saline
BrdU: 5-Bromo-2’-deoxyuridine
mJ: Millijoule
TBS: Tris-buffered saline
DAPI: 4’,6-diamidino-2-phenylindole
GFP: Green fluorescent protein
DsRed: Red fluorescent protein
Cy5: Cyanine 5
T-T dimer: Thymine dimer
CC3: Cleaved caspase-3

---

**Supplemental Figure 1.**
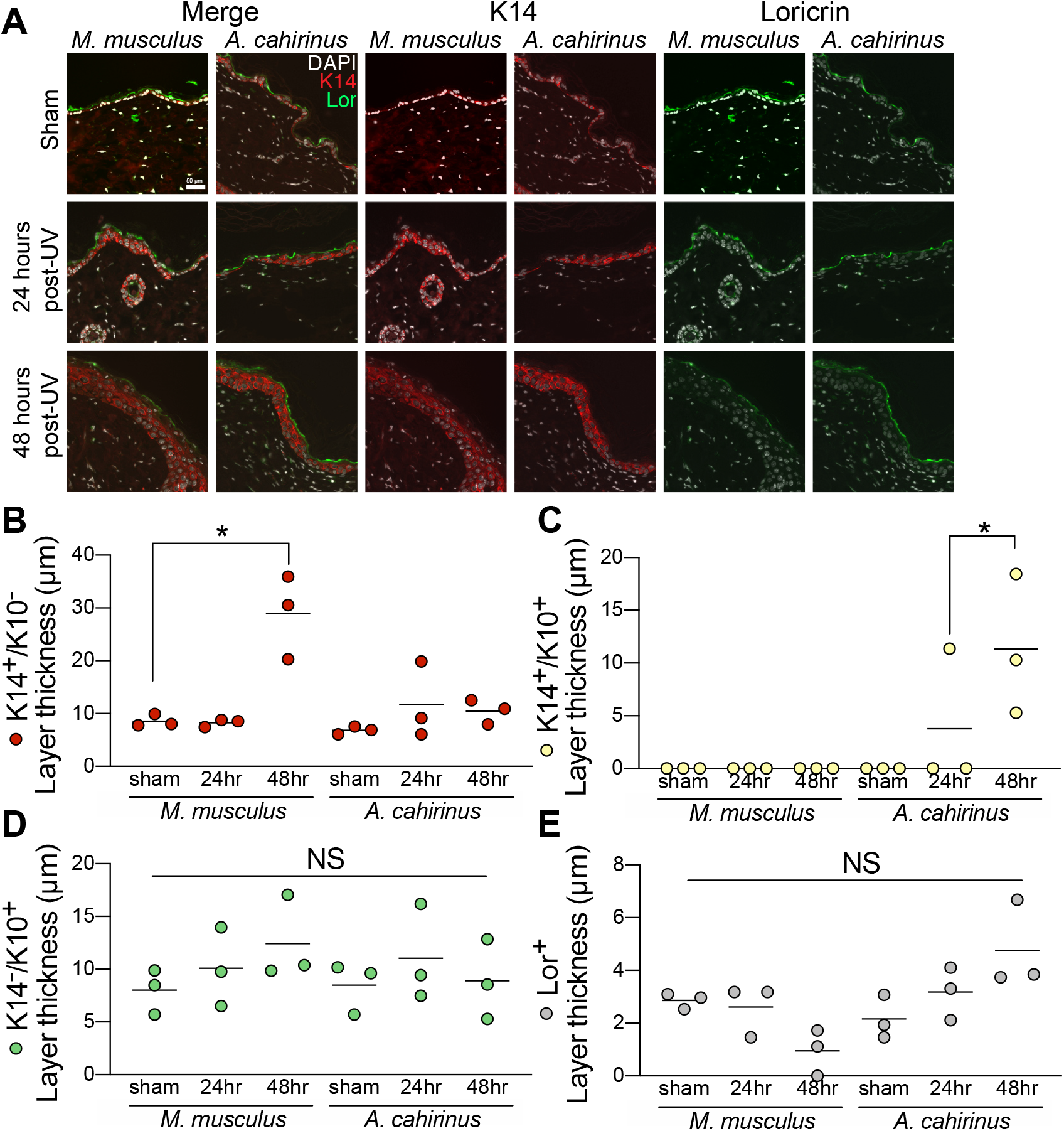
Acute UVB-exposure induces distinctive pattern of skin epidermal differentiation in *A. cahirinus*. *A*, representative immunofluorescence images of epidermal differentiation markers keratin 14 (*K14*) and loricrin (*Lor*) labeling with magnified inset of the basal-suprabasal junction. *B, C, D, E* individual layer thickness quantification of the (*B), K14*^+^/*K10*^−^ single positive basal layer *(C), K14*^+^/*K10*^+^ double positive middle suprabasal layer *(D), K14*^−^/*K10*^−^ single positive spinous layer and *(E), Lor*^+^ single positive cornified envelope. *n = 3 mice per group.* Data points are biological replicates. Lines indicate group means. *, significantly different from the indicated group (*p<0.05*).

